# Characterisation of factors contributing to the performance of nonwoven fibrous matrices as substrates for adenovirus vectored vaccine stabilisation

**DOI:** 10.1101/2020.01.31.928218

**Authors:** Pawan Dulal, Adam A Walters, Nicholas Hawkins, Tim DW Claridge, Katarzyna Kowal, Steven Neill, Stephen J Russell, Adam Ritchie, Rebecca Ashfield, Adrian VS Hill, Alexander D Douglas

## Abstract

The global network of fridges and freezers known as the “cold chain” can account for a significant proportion of the total cost of vaccination and is susceptible to failure. Cost-efficient techniques to enhance stability of vaccines could prevent such losses and improve vaccination coverage, particularly in low income countries. We have previously reported a novel, potentially less expensive thermostabilisation approach using a combination of simple sugars and glass micro-fibrous matrix, achieving an excellent recovery of vaccines after storage at supraphysiological temperatures. This matrix is, however, prone to fragmentation and currently not suitable for clinical translation.

Here, we report an investigation of alternative, potentially GMP compatible, fibrous matrices. A number of commercially-available matrices permitted good protein recovery, quality of sugar glass and moisture content of the dried product but did not achieve the thermostabilisation performance of the original glass fibre matrix. We therefore further investigated physical and chemical characteristics of the glass fibre matrix and its components. Our investigation shows that the polyvinyl alcohol present in the glass fibre matrix assists vaccine stability. This finding enabled us to develop a custom-produced matrix with encouraging performance, as an initial step towards a biocompatible matrix for clinical translation. We discuss the path to transfer of the technology into clinical use, including potential obstacles.

## Introduction

Vaccine immunogens are composed of complex biological macromolecules. Good immunogenicity requires stability of these molecules throughout the lifespan of a product, from production and formulation, through transportation and storage to delivery to the recipient. Extrinsic factors such as light, pH, agitation and oxidants combine with temperature fluctuations to challenge product stability. Therefore, most vaccines need to be continuously stored in fridges or freezers. Maintaining the cold chain and associated logistics in vaccination campaigns can contribute up to 45% of the total cost of vaccination ^1^. On the other hand, damage as a result of cold chain breakages costs several million dollars annually ^2^. These issues affect not only low income countries, where the cold chain is regarded as being least reliable, but also developed countries ^3 4 5^. The vaccines responsible for eradication of smallpox and rinderpest (the only two diseases eradicated by immunisation) were both thermostable, a factor believed to contribute to their success ^6^. Development of technologies to enhance vaccine thermostability has therefore been a major focus of research effort ^7^.

Viral vectored vaccines, in particular adenoviruses, are versatile platforms for development of novel vaccines against malaria ^8 9^, HIV-AIDS ^10 11^, tuberculosis ^12^ and influenza ^13 14^. They have provided one of the leading approaches to control of the recent Ebola epidemics, and the speed of producing new vaccines using a generic manufacturing process makes them attractive for emergency response to other outbreak pathogens ^15, 16^. Successful thermostabilisation of adenovirus vectored vaccines could thus be valuable for products targeting a wide range of diseases.

There have been a number of efforts to enhance stability of adenovirus-based vaccines in liquid formulation ^17–19^. Other groups have reported improvement in stability by drying vaccines using lyophilisation ^20, 21^, spray drying ^22–24^, and nano-patch technology ^25^. We have previously reported that drying a formulation of adenovirus in glass forming sugars upon a fibrous matrix such as paper or a wetlaid nonwoven fabric permitted full recovery of viable and immunogenic virus, even after six months at 45°C ^26–28^. We refer to this method as sugar-matrix thermostabilisation (SMT). As well as thermostabilisation performance, advantages of SMT include a relatively short process duration (in some cases as little as 18 hours) and avoidance of the stress of extreme in-process temperature exposure involved in lyophilisation and spray drying.

Our previous published SMT work used a glass fibre matrix and a polypropylene matrix; we reported that the glass fibre matrix achieved better stability than the polypropylene matrix ^26^. Glass fibre matrix is however prone to shedding of non-biocompatible fibres, and so not suitable for clinical translation. Here, we present results from an effort to characterise the factors contributing to the performance of glass fibre matrix and to test alternative matrices more suitable for clinical use.

## Methods

### Viruses and infectivity titration

Simian adeno virus vectors ChAd63-METRAP and ChAdOx2-RabGP were prepared, purified and tested for quality by the Jenner Institute Viral Vector Core Facility, as previously described ^28, 29^. Viruses were dialysed against either a previously used storage buffer (10mM Tris, 7.5 % w/v sucrose, pH 7.8) or unbuffered 0.5M trehalose and sucrose and stored at −80°C as stock. Typical preparations were supplied and stored at a titre of c. 1×10^12^ virus particles (VP) per mL, corresponding to c. 1×10^10^ infectious units (IU) per mL and hence a particle: infectivity ratio of c. 100.

For infectivity titration, duplicate fivefold serial dilutions were prepared in complete DMEM (10% FCS, 100 U penicillin, 0.1 mg streptomycin/ml, 4 mM L-glutamine) and used to infect 80-100 % confluent HEK293-TRex cells (ThermoFisher) grown in 96-well plates (BD Purecoat Amine, BD Biosciences, Europe). Infected cells were immunostained and imaged as previously described ^30^. Wells containing 20-200 spots were used to back-calculate recovered infectious units.

### Drying, thermochallenge and reconstitution

Stock vaccines were thawed and diluted into unbuffered 0.5M trehalose – sucrose solution. Dilution factors, final viral titres, and the trehalose: sucrose ratio varied between experiments, as stated in figure legends.

Fibrous matrix was cut into 100 mm^2^ pieces, loaded with vaccine formulation and transferred to a glove box (Coy Laboratory Products). An activated silica bed within the chamber and circulation of air through desiccation capsules containing anhydrous calcium sulphate (Drierite^TM^, W.A. Hammond Drierite Co.), regulated by a humidity controller (Series 5000, Electro-tech Systems), was used to maintain relative humidity beneath 5%. A portable datalogger (AET-175, ATP instruments) was used for recording changes in relative humidity and temperature during the desiccation process. The temperature inside the enclosed glove box remained between 22°C and 25°C for all experiments.

Samples were transferred into 2 mL glass vials, stoppered and crimp sealed under dry conditions within the glove box prior to further use. Samples undergoing thermochallenge i.e. storage at elevated temperature (typically 45°C for one or four weeks) were stored within secondary packaging (moisture barrier bags).

For experiments involving reconstitution of dried samples, this was performed by addition of phosphate buffered saline (Sigma), followed by brief vortexing of the vial (1±0.5 seconds, three times). Virus infectivity after reconstitution was assayed as described above. Recovery of infectious virus was quantified by comparison to a sample of the starting material, included on the same assay plate. Between the set-up of an experiment and the assay of recovered infectivity, such comparator material was stored at −80°C in aqueous buffer (under which conditions loss of infectivity is known to be negligible).

Recovery was calculated in terms of log_10_-fold loss in the total infectious virus content of the matrix i.e. log_10_-fold loss = log_10_(infectious units dried on matrix based on −80 stored sample) – log_10_(infectious units recovered from matrix). Unlike viral titres (infectious units / mL), this parameter is independent of the volumes in which sample was dried or reconstituted. 0.3 log_10_-fold loss thus implies c. 50% recovery, 0.5 log_10_-fold loss implies c. 30% recovery, and 1 log10-fold loss implies 10% recovery.

### Karl Fischer moisture analysis

Residual moisture content in single matrix post-desiccation was determined with a Karl Fischer moisture analyser equipped with coulometer (Metrohm), and Hydranal-Coulomat titration solution (Honeywell, Fluka) in accordance with the manufacturers’ recommendations. A standard was used to calibrate the instrument performance (lactose standard 5%, MerckMillipore). Residual moisture content was expressed as the mass extracted per 50 µL of formulation loaded into each matrix.

### Measurement of subvisible particles

100 mm^2^ pieces of glass fibre (S14) were cut, autoclaved at (121 °C, 15 minutes) and loaded with 50 µL of sugar solution prior to drying in the glove box, vialling and reconstitution as described above. Reconstituted solution from 10 vials was aspirated using a syringe with a conventional 20G needle (BD Biosciences), to produce a single pooled unfiltered sample. Reconstituted solution from a further 10 vials was aspirated using a 5-micron filter needle (BD Biosciences), to produce a single pooled filtered sample. Both pools were diluted to a final volume of 25 mL and tested for sub-visible particles by light microscopy according to Ph.Eur (2.9.19) / USP <788> by a commercial testing laboratory (Reading Scientific Services Limited).

### Scanning electron microscopy

Untreated matrices (i.e. without sputter coating) were loaded onto aluminium mounts using carbon conductive tabs and imaged using a Zeiss-Evo LS15 variable pressure scanning electron microscope (SEM) equipped with variable pressure secondary electron detector (Carl Zeiss Ltd). Imaging was performed at a chamber pressure of 50 Pa air and accelerating voltage of 15 kV. Further analyses of images such as measurement of fibre diameter were made using Image J software. At least 40 measurements per matrix were performed, using images taken at varying magnification.

### Differential scanning calorimetry

The glass transition temperature (T_g_) of sugar glass in matrices was measured immediately after drying had completed using Differential Scanning Calorimetry (DSC). The melting point of the binder in untreated glass fibre (S14) was measured using DSC (Q2000, TA instruments). The instrument was purged with dry nitrogen (50 mg/mL) continuously during sample measurement. Calibration was performed prior to measurements using a certified reference material (Indium) for temperature and heat flow accuracy.

Multiple 6-millimetre diameter discs were cut out of a matrix dried with 0.5M trehalose: sucrose (50:50). Discs were weighed and, for each matrix type in turn, a total mass of 5-15 milligrams was loaded into Aluminium DSC pans (TA Instruments) and hermetically sealed. Samples were subjected to a temperature ramp from −20 °C to 180 °C at a heating rate of 10 °C per minute. Measurements on all samples were performed in duplicates. Thermograms relating heat flow (W/g) to temperature (°C) were analysed using Trios software (TA Instruments) for identification of the glass transition (Tg) onset temperature.

For the measurement of enthalpic recovery, which manifests as an endothermic peak at the glass transition, modulated DSC (temperature modulation ±0.50°C every 60 seconds and ramp rate 3°C/min from −20°C to 100°C) was employed. The enthalpic recovery was estimated by linear peak integration in the thermograms plotted between nonreverse heat flow (W/g) and temperature (°C) using Universal Analysis software (TA Instruments).

### Protein recovery

A 10mg/mL solution of lysozyme (Sigma-Aldrich) was made in 0.5M trehalose sucrose. 25 µL was loaded into each matrix in triplicates. Protein was reconstituted from the matrices after desiccation overnight and recovery was quantified using EnzChek Lysozyme Assay Kit (ThermoFisher Scientific). Fluorescence measurements were performed in triplicates for each sample and protein recovery calculated by interpolation on a standard curve, using GraphPad Prism.

### Thermogravimetric analysis

Degradation temperatures of matrix constituents were measured by thermogravimetric analysis using a TGA Q500 (TA instruments). Samples loaded into a tared platinum pan just prior to measurement were subjected to a temperature ramp at 5°C per minute from ambient to 550°C in a flowing nitrogen atmosphere (100ml/min). The gas was switched to air at 550°C (100 ml/min) and heat was continued at the rate of 5°C/min to 730°C. Data was analysed using Universal Analysis software (TA Instruments).

### Polyvinyl alcohol (PVA) extraction and Fourier transform infrared spectroscopy

A 500 mm X 27 mm piece of glass fibre (S14) was dissolved in 100 mL of ultrapure water by stirring and heating at 90 °C for 48 hours. The suspension was filtered through a 0.2 µm filter and the filtrate was freeze dried (Virtis AdVantage 2.0, SP Scientific) to a white amorphous material. This residue was subjected to single reflection attenuated total reflection (MIRAacle ^TM^, Pike Technologies) Fourier transform infrared spectroscopy ATR-FTIR (Tensor 37, Bruker) equipped with nitrogen-cooled mercury cadmium telluride detector.

To obtain a spectrum of the sample, an average of 64 interferograms was collected at a resolution of 4 cm^-1^ in the wavelength range from 4000 cm^-1^ to 750 cm^-1^ and blank subtracted.

Spectra were analysed using OPUS 6.5 software (Bruker) and compared with RMIT University’s spectral library of organic compounds generated using Spectrum 10 software (Perkin Elmer). Fingerprint spectra shown in Figure 4D were prepared using GraphPad Prism.

### Dynamic light scattering (DLS)

The molecular weight of the PVA binder present on the glass fibre matrix (S14) was estimated by DLS. PVA was extracted and freeze-dried as described above. Solutions of this extracted sample and standards of known polymer size were prepared in water to the concentration of approximately 1.3 mg/mL and passed through 0.22 µm and 300 kDa molecular weight cut-off filters. The sample was then concentrated approximately five-fold from 4mL to 0.75 mL using a 3KDa MWCO filter (Vivaspin, GE). Zetasizer Nano ZS and DTS software (Malvern Instruments) was used for measurement of hydrodynamic diameter based on size distribution by volume. Independent duplicate preparations of all standards and samples were tested. GraphPad Prism was used to generate a standard curve plotted between measured hydrodynamic diameter and known average molecular weight (KDa) to interpolate size of the PVA in the extract.

### Nuclear magnetic resonance (NMR) spectroscopy

The degree of hydrolysis of the PVA binder present on the glass fibre matrix (S14) was estimated by ^1^H NMR measurement as the intensity of the peak attributable to the acetyl group present in non-hydrolysed PVA. A 5 mg/mL aqueous solution of PVA extracted from the glass fibre sample (S14) was prepared in deuterium oxide. Reference spectra for PVA with varying degrees of hydrolysis were obtained by mixing >99% hydrolysed PVA and 80% hydrolysed PVA in appropriate proportions to produce standards containing c. 0%, 2%, 5%, 10% and 20% acetyl groups, again at 5 mg/mL in deuterium oxide. An AVIII 700 instrument (Bruker Biospin) was used to generate ^1^H 1D spectra (employing a quantitative 1D NOESY (Nuclear Overhauser Effect Spectroscopy) presaturation sequence with recovery delay d_1_ =30s) and 2D ^1^H-^13^C HSQC (heteronuclear single quantum coherence spectroscopy) plots.

### Application of PVA to matrices

Aqueous solutions of each type of PVA to be investigated were prepared at 10 mg/mL. Matrices were cut into approximately 100 mm^2^ pieces and loaded with until the matrices were saturated by the solution and air-dried overnight. The polyamide matrix (33100L) required surface modification by washing in 100% ethanol for 20 minutes and air drying before PVA could be loaded. Following PVA application, vaccines were dried on the matrices and thermostability assessed as described above.

### Study of physical characteristics of matrices

The physical characteristics of matrices were studied using NWSPs (Nonwoven Standard Procedures) prescribed by the EDANA, the international trade association for the nonwoven industry.

Areal density was measured as ratio of mass in grams and area in square metres according to EDANA NWSP 130.1R0 (15).

Matrix thickness was measured at an applied pressure of 0.5 kPa according to EDANA NWSP 120.1.R0 (15) using a thickness tester (ProGage, Thwing-Albert Instrument Company).

Fibre length was measured based on 60 individual fibres extracted from the matrix using a Leica MD G41 optical microscope in transmission mode and Image J software.

Minimum mean and maximum pore size were obtained using a POROLUX 100 Automated Capillary Flow Porometer using Galpore liquid of surface tension 15.6 mN·m^-1^. A total of five measurements were taken per sample. Porosity (ɛ) of the GE S-14 was determined by the equation ɛ= (1−Ø) X 100 % where Ø is the volume fraction measured as a ratio of bulk matrix density to bulk fibre density.

The surface tension of aqueous sugar solution on the matrix was measured on a Kruss K100 tensiometer using the Wilhelmy plate method in which the force exerted on a suspended plate when it touches the surface of a liquid is related to the surface tension and the contact angle according to the equation 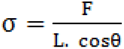 where σ= surface tension of the liquid, F=measured force, L=wetted length, and Θ=contact angle.

Wettability was evaluated using the Washburn method, again on a Kruss K100 tensiometer. The rate of mass uptake when the porous substrate (matrix) comes in to contact with a liquid was used to determine the capillary constant of the substrate by applying the Washburn equation 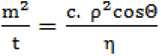, where m=mass, t=flow time, c=capillary constant of the nonwoven, ρ=density of the liquid, σ= surface tension of the liquid, Θ=contact angle, η=viscosity of the liquid. The capillary constant of the matrix was determined using n-hexane which has a contact angle of 0°.

### Development of Nonwoven Fabrics for Vaccine Thermostabilisation

A glass fibre matrix (Figure 5B) was custom made by a wet-lay process using commercially available glass fibre and polyvinyl alcohol aiming for similar porosity, thickness, areal density and wetting behaviour as the glass fibre sample (S14). A matrix of glass fibre type 475 with diameter 4 µm (Johns Manville) was prepared and 1 g/L PVA solution (>99% hydrolysed, Mw 146000-186000 kDa, Sigma) was applied to each side using a spray gun before drying at 110°C for 15 minutes.

## Results

### Glass fibre matrix achieves good stability but is not suitable for clinical development

In our previously published work, we had used low vaccine doses (1.1 × 10^10^ viral particles per matrix, as compared to a typical human dose of 5 x 10^10^ viral particles), applied to the glass fibre matrix ‘Standard 14’ (S14, GE Healthcare, Figure 1A) ^26^. We speculate that the fibrous matrix provides a high surface area, favouring relatively rapid drying of the product despite the gentle conditions used (18-24 hours at 20-23 °C and atmospheric pressure). The process results in films of vitreous ‘sugar glass’ embedded in the matrix (Figure 1B). As previously reported, this ‘base-case’ process achieves a sugar glass transition temperature (T_g_) after drying of 47-55°C, with moisture content 3-5% of total solute dry weight ^26^.

**Figure 1:**
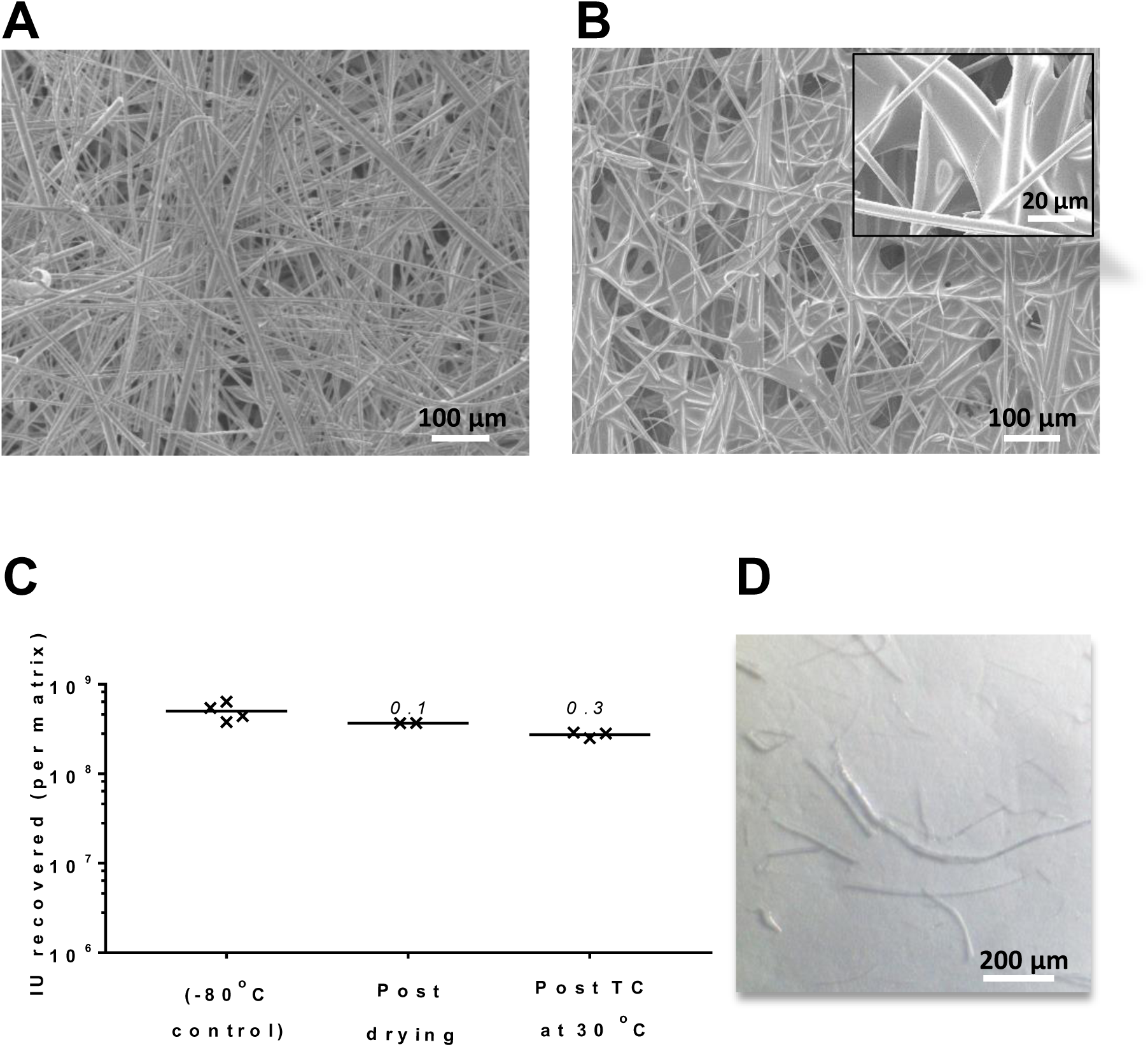
SMT on S14 results in thermostable vaccine embedded in sugar glass, Panels A and B show scanning electron micrographs of S14, respectively before and after loading and drying sugar formulation. In Panel A, the arrangement of fibres in S14 is apparent (200X magnification, 100 µm scale bar). In panel B, sugar glass films are visible between fibres (main panel at 200X magnification with 100µm scale bar, inset at 1000X magnification with 20µm scale bar). Panel C shows recovery of vaccine from matrix loaded with a human dose of adenovirus, both immediately post-drying and after thermochallenge at 30 °C for one month. Results plotted illustrate total virus recovery from each replicate matrix (n=4 for −80 °C comparator, n=2 for post-drying, n=3 for post-thermochallenge), with lines indicating the mean of the replicates. Numbers above each result show log10-fold loss from the −80 °C control. Panel D shows a representative light microscopy image of fibres shed from S14 after vaccine reconstitution (scale bar 200 µm).

We have now dried full human doses (5×10^10^ virus particles, approximately 5×10^8^ infectious units [IU]) of a simian adenovirus vectored rabies vaccine (ChAdOx2 RabG) upon 1 cm^2^ of the matrix, with <0.1 log_10_-fold loss in-process (i.e. during desiccation) and approximately 0.3 log_10_-fold loss after thermochallenge at 30°C for a month (Figure 1C) ^28^. For explanation of the log_10_-fold loss metric, please see Methods.

Despite the good thermostabilisation performance of the glass fibre matrix, concerns were raised regarding the brittleness of glass fibre would result in shedding of fibres during the process of vaccine reconstitution. Indeed, macroscopic damage to the matrix was sometimes apparent at the point of reconstitution. We therefore sought to quantify subvisible particles (0.1-100 micron) in the reconstituted vaccine. Application of a pharmacopoeial light microscopy method revealed numerous glass fibres (Figure 1D). Although passing the reconstituted solution through a 5 µm filter needle removed virtually all detectable glass particles, reducing the particulate burden within pharmacopoeial limits, such a method is unsuitable for clinical development, with the possible exception of very early-phase studies ^31^.

### Selection of commercially-available matrices for evaluation

In our previously published work, we reported that the S14 matrix achieved better vaccine thermostability than an alternative commercially-available polypropylene-based matrix (HDC^®^II J200, Pall Corporation). We therefore sought to identify alternative commercially-available matrices which might offer thermostability equivalent to or better than that achieved with S14, but without the problem of shedding of non-biocompatible fibres. A set of seven matrices were selected based on manufacturers’ product specifications claiming low fibre shedding, low chemical leaching and compatibility with sterilisation either by dry heat, steam or gamma-radiation sterilisation (Table 1). Henceforth matrices are referred to, for clarity of identification, in terms of their fibre material and the manufacturer’s product name.

**Table 1:** Basic characteristics of studied matrices

| Matrix name<br>(manufacturer product<br>reference) | Manufacturer (sub-brand if<br>applicable) | Main material | PS20 treatment? | Absorption >20 $\mu\text{L}/\text{cm}^2$ $\text{H}_2\text{O}$ | Loading capacity<br>($\mu\text{L}/\text{cm}^2$) | Fibre diameter<br>($\mu\text{m}$) ±SEM | Mean pore size<br>($\mu\text{m}$) | Thermostabilisation<br>performance ($\log_{10}$-fold<br>infectivity loss) |
| --- | --- | --- | --- | --- | --- | --- | --- | --- |
| S 14 | GE<br>(Whatman) | Glass fibre | X | ✓ | 55 | 4.2±0.3 | 20 | 0.3 |
| J200 | Pall | Polypropylene | ✓ | ✓ | 36 | 21.7±0.4 | 20 | 1 |
| 23100 | Hollingsworth<br>- Vose | Polyester | ✓ | ✓ | 43 | 4.4±0.4 | 11 | 1.3 |
| 33100L | Hollingsworth<br>- Vose | Polyamide | ✓ | ✓ | 36 | 4.8±0.4 | 27 | 0.9 |
| Cyclopore<br>(PC) | GE<br>(Whatman) | Polycarbonate | ✓ | X | Excl. | Excl. | Excl. | Excl. |
| 31 ET CHR | GE<br>(Whatman) | Cellulose | X | ✓ | 40 | 15.4±1.0 | 1.2 | 2.4 |
| Leukosorb<br>(Leu) | Pall | Polyester<sup>a</sup> | X | ✓ | 40 | 3.5±0.2 | 8 | 1.7 |
| Conjugate<br>pad (CP) | Pall | Glass fibre<sup>b</sup> | X | ✓ | 30 | 12.3±0.4 | N/A | 0.9 |
| Asymmetric<br>polysulfone<br>(AS) | Pall | Polysulfone | ✓ | X | Excl. | Excl. | Excl. | Excl. |

Loading capacity was measured by weight measurement of matrices before and after full impregnation in water. Fibre diameter was measured using Image J from SEM micrograph images. Mean pore size are as provided in suppliers' specification (in one case N/A denotes data not available). Thermostabilisation performance presented is loss of infectivity after vaccine drying and thermochallenge for 1 week at 45 °C. Excl. denotes matrices excluded from further analysis on basis of loading capacity.
<sup>a</sup> Proprietary composition, but macroscopic, microscopic and FTIR spectrum appearances are consistent with polyester (data not shown); this matrix is henceforth referred to as polyester
<sup>b</sup> Henceforth referred to as 'glass fibre (conjugate pad)' for avoidance of confusion with S14.

We initially screened all matrices for suitable loading capacity (>20 µL of deionised water/cm^2^). Matrices which did not absorb 20 µL/cm^2^ (without visible remaining beads of water within two seconds) were re-tested after treatment with 2% polysorbate 20 solution. Matrices which did not absorb 20 µL/cm^2^ after detergent treatment were not studied further. We proceeded to further study of the remaining five matrices along with the two previously tested matrices (glass fibre S14 and polypropylene J200).

The architecture of the selected matrices was characterised by scanning electron microscopy (Figure 2). Fibre diameters estimated for each matrix type using Image J analysis are presented in table 1.

**Figure 2:**
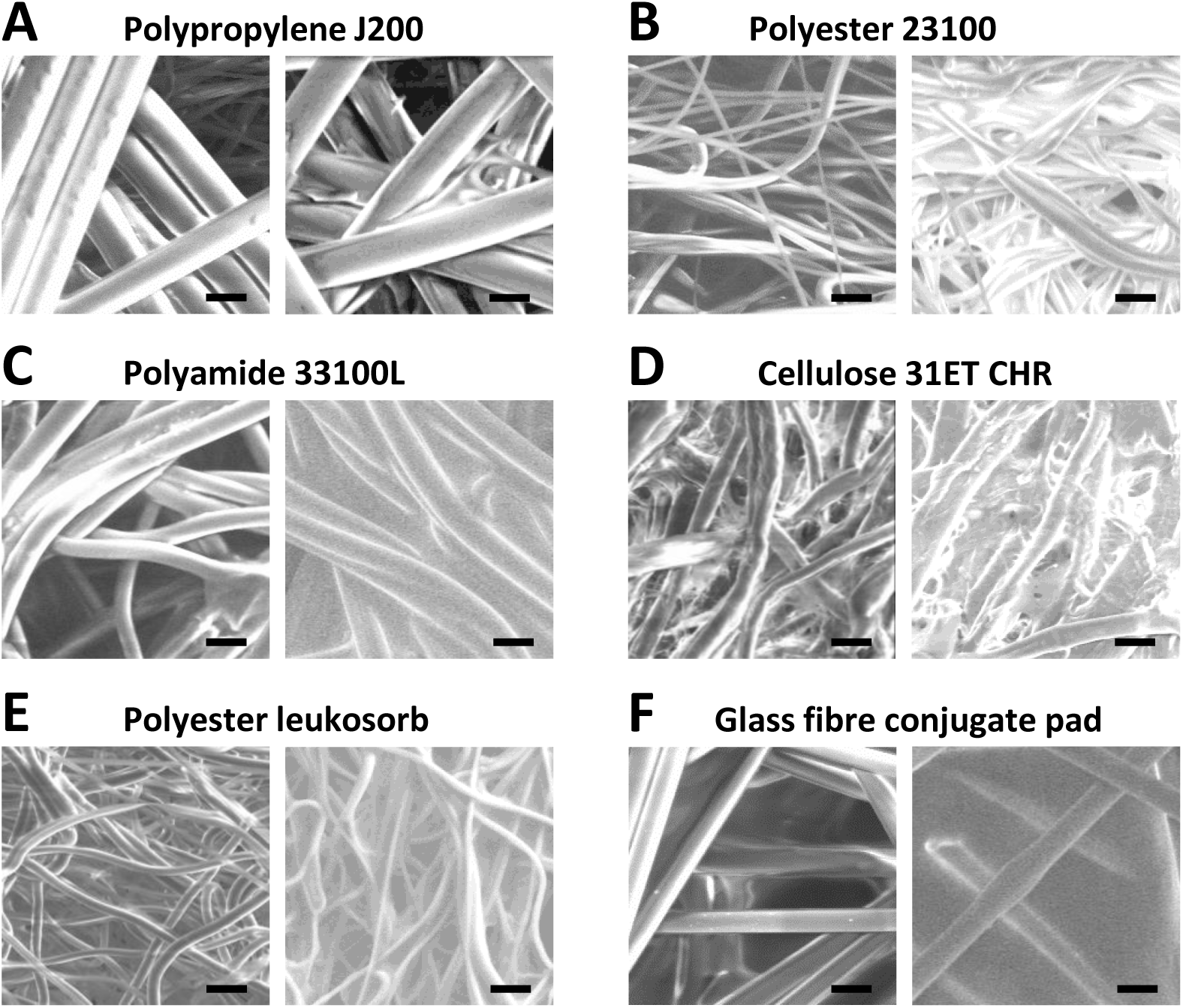
Physical appearance of fibres and sugar glass formed in matrices. Scanning electron microscopy images of six different matrices at 1000X magnification. Within each panel, the left-hand image shows empty matrices and the right-hand image shows sugar-loaded fibres. Scale bar shows 20 µm in each image.

Fibres in the polypropylene (J200) matrix (Fig 2A) and glass fibre (conjugate pad) matrix (Fig 2F) shared the straight, rod-like fibre morphology seen in the original glass fibre (S14) matrix (Fig 1A-B) but had larger fibre diameters of 10 – 20 μm (as compared to 4 μm in the S14 glass fibre sample). The remaining four matrices (Fig 2B-E) exhibited a greater degree of curl along their length. A film, potentially a binder, was apparent on the glass fibre matrix (conjugate pad) (Fig 2F). Matrices loaded with 0.5 M sugar solution and dried at room temperature for 24 hours also showed differences in the distribution of the sugar glass intercalated between the fibres**Error! Reference source not found.**(Figure 2). Distinct films of sugar glass between the fibres were visible on the two glass fibre matrices, but not in the polyamide (33100L), polyester (leukosorb) or polypropylene (J200) matrices. The sugar loaded polyester (23100) matrix and cellulose (31 ET CHR) matrices showed a glazing effect on the fibres, with discrete sugar glass films being apparent.

### Thermostability of adenovirus vaccine formulated in commercially-available matrices

Adenovirus vaccine vectors were formulated in 0.5M TS and dried on the five selected ‘new’ matrices, with glass fibre (S14) and polypropylene (J200) matrices as comparators of known performance. Dried matrices were thermochallenged for a week at 45°C prior to reconstitution and infectivity titration. Marked thermostabilisation performance differences between the matrices were apparent (Table 1).

As previously observed, the glass-fibre matrix (S14) showed minimal (less than 0.3 log_10_-fold) loss of infectivity compared to the −80 °C stored positive control. None of the alternative matrices matched this level of performance: the next-best-performing matrices were polyamide (33100L) and glass fibre (conjugate pad), with infectivity loss of 0.9 log_10_-fold in each. The cellulose-based matrix (GE) performed most poorly.

### Relationships between matrix type, thermostabilisation performance and physical characteristics of sugar glass

In order to understand the highly variable thermostabilisation performance of the matrices, we investigated whether performance could be correlated with the physical properties of the matrices or the sugar glass formed.

Standard sugar formulation (0.5M trehalose sucrose) was loaded into each matrix and dried at room temperature and < 5% relative humidity. Dried samples were then subjected to modulated differential scanning calorimetry (DSC) to measure glass transition temperature (Tg) onset temperature and enthalpic recovery of the sugar glass formed on the matrices. The glass transition temperature (T_g_) indicates the temperature at which a low mobility sugar glass changes to a highly mobile rubbery state and is known to be related to product stability in dry formulations ^32^. Enthalpic recovery is a measure of energy dissipated as a glass progresses through equilibrium and can reflect molecular rearrangement during storage or physical ageing ^33^.

The T_g_ was similar for all matrices, observed over a narrow range between 52°C and 56°C, and did not correlate with thermostability (Figure 3A). A possible correlation was observed between thermostabilisation performance and high enthalpic recovery (Figure 3B, r^2^=−0.70, p=0.02), as seen with glass fibre matrices conjugate pad and S14 followed by polypropylene (J200) based matrix. There was substantial variation in the residual moisture content of products dried under the same conditions on different matrices (Figure 3C), with lowest residual moisture in the best performing matrix, S14. Recovery of a model protein (lysozyme) after desiccation and reconstitution did not predict thermostabilisation performance (Figure 3D).

**Figure 3:**
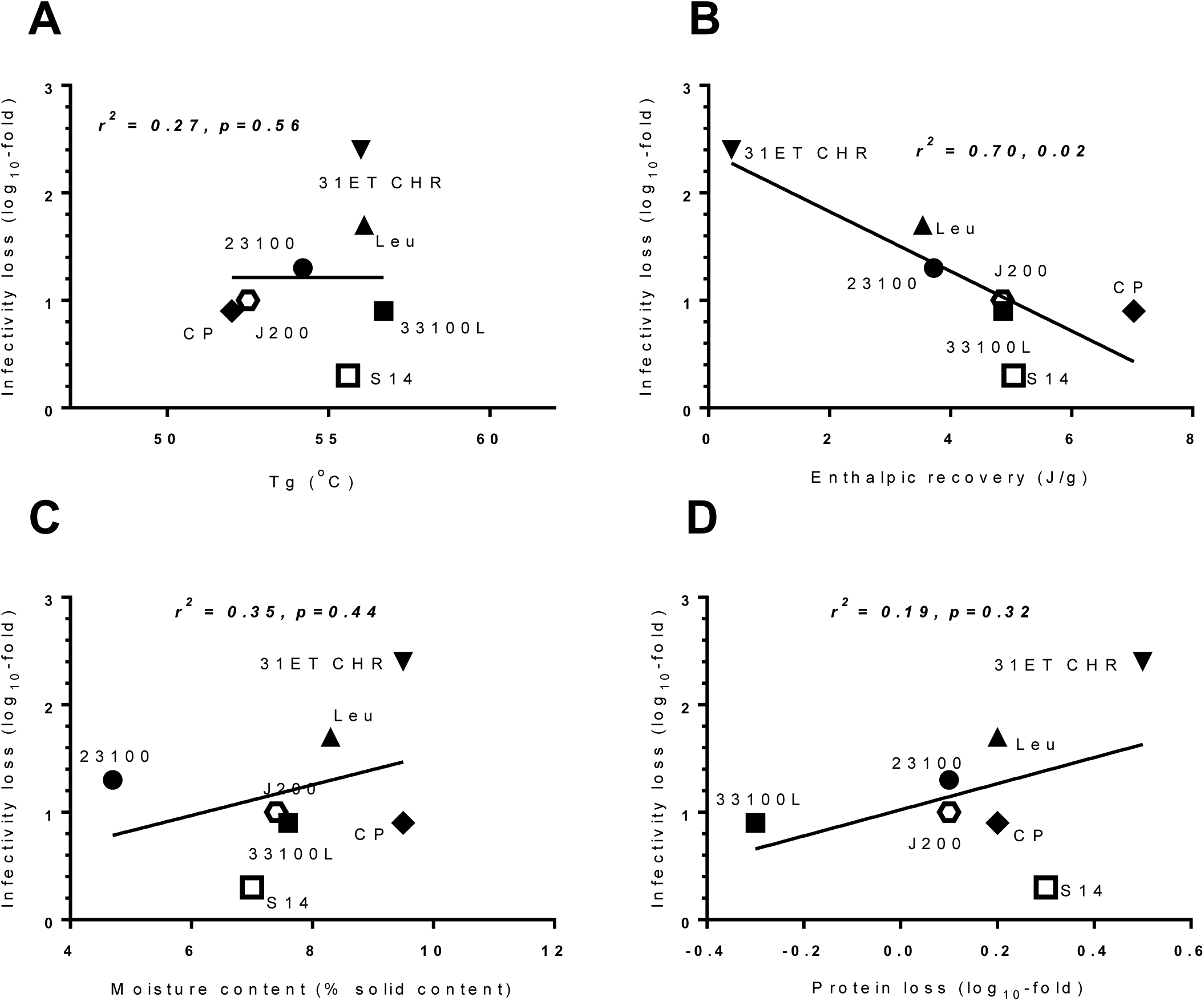
Physical properties of sugar glass have differential impact on vaccine thermostability. Each panel relates the thermostabilisation performance of the various matrices (after one-week thermochallenge at 45 °C, Y-axis, data shown in Table 1) to a potentially explanatory variable on the X-axis: glass transition temperature (Panel A); enthalpic recovery (Panel B); moisture content (Panel C); and protein recovery (Panel D). Results of Pearson correlation analysis are shown within each panel. DSC measurements (Panels A and B) were made in singlicate, Karl-Fischer measurements (Panel C) were made in duplicate, and protein recovery measurements (Panel D) were made in triplicate. For panels C and D, points represent the mean measurement.

### Characterisation of S14 glass fibre matrix

Given our inability to identify a commercially-available matrix suitable for clinical application, we turned our attention to detailed characterisation of the best-performing S14 matrix, with a view to future production of a similar matrix from biocompatible materials.

We initially performed a more extensive characterisation of the physical properties of the matrix, with results as shown in Table 2.

**Table 2:** Physical characteristics of the glass fibre (S14) matrix

| PARAMETER (unit) | Mean | Standard deviation |
| --- | --- | --- |
| Area density [$\text{g.m}^{-2}$] | 55.1 | 1.8 |
| Thickness [mm] | 0.5 | 0.04 |
| Fibre diameter [$\mu\text{m}$] | 3.4 | 1.8 |
| Fibre length [mm] | 4.4 | 1.5 |
| Largest pore diameter [$\mu\text{m}$] | 133.0 | 99.4 |
| Mean flow pore diameter [$\mu\text{m}$] | 22.3 | 0.5 |
| Smallest pore diameter [$\mu\text{m}$] | 6.4 | 1.5 |
| Porosity [%] | 95 |  |
| Capillary constant [$\text{cm}^5$] | $7.7 \times 10^{-7}$ | $5.6 \times 10^{-8}$ |
| Thermostabilisation buffer contact angle [$^\circ$] | 30.9 | 4.9 |
| Thermostabilisation buffer surface tension [$\text{mN.m}^{-1}$] | 63.9 | 0.3 |

Use of chemical binders is common in the production of nonwoven fabrics such as S14 to provide strength and desirable surface properties. In view of the possibility that a binder might be contributing to S14’s thermostabilisation performance, we investigated whether such a binder could be identified in the matrix.

Scanning electron microscopy demonstrated film-like material which could represent binder covering fires in some sections of the S14 sample (Figure 4A). Differential scanning calorimetry and thermogravimetric analysis (Figures 4B-C) demonstrated the presence of a material with a melting point of 220°C and a degradation temperature at 260°C.

**Figure 4:**
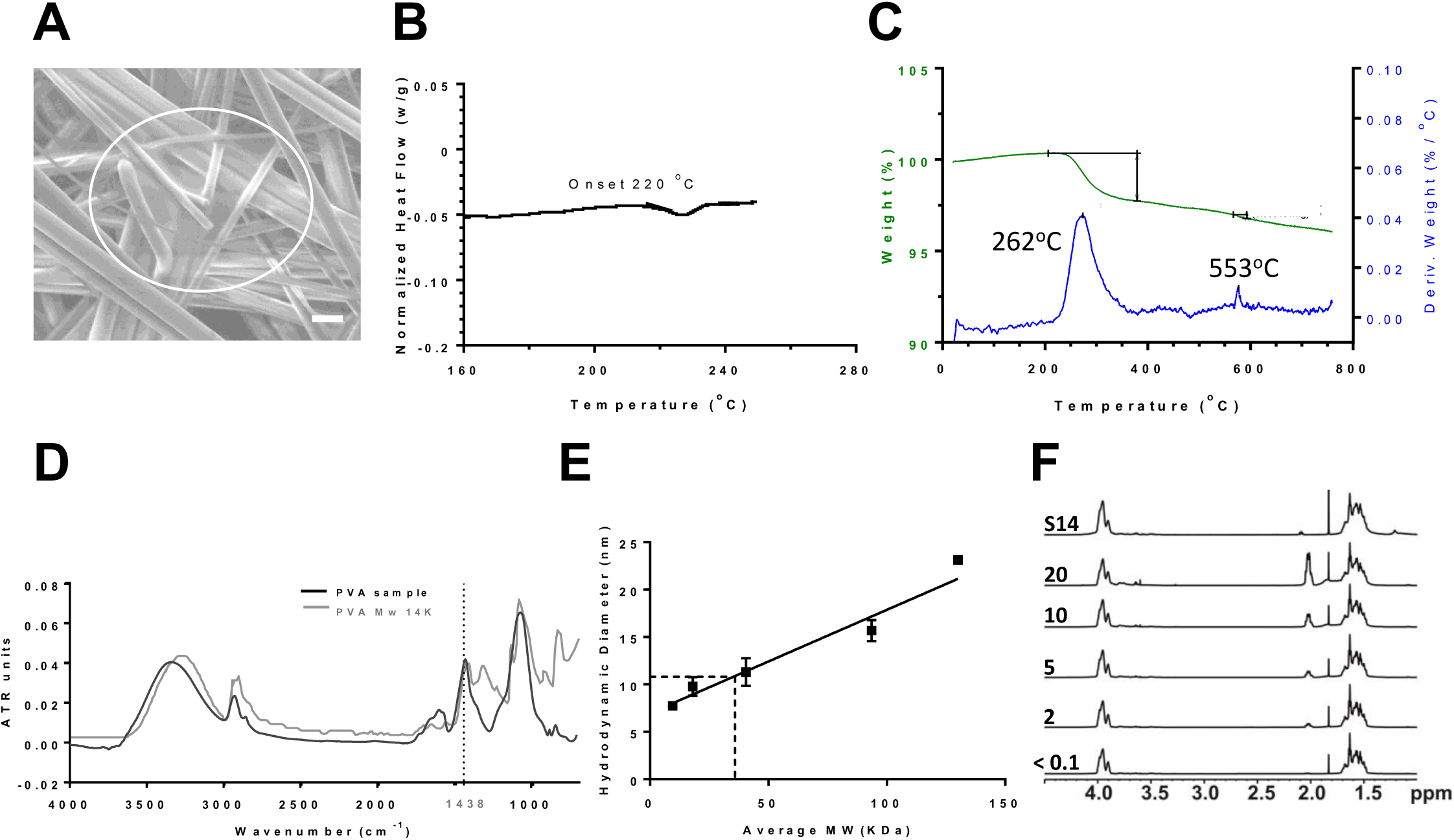
Chemical properties of glass fibre matrix. Panel A shows a scanning electron microscopy image indicating presence of binder in glass fibre (S14), visible as a film in the white-circled area. Scale bar shows 10 µm in each image. Such areas were relatively sparse, compared to the extensive films apparent in sugar-loaded matrices (Figure 1B). Panel B shows a differential scanning calorimetry thermogram of glass fibre (S14), representative of duplicate measurements. The graphs show heat flow as a function of temperature during scanning from 40 °C to 250 °C. Panel C displays thermogravimetric analysis of glass fibre (S14), showing a thermal decomposition step. Weight losses (green lines) and the rate of weight loss (i.e. derivative, %/°C) (blue lines) are shown. Panel D shows the sample spectrum obtained from attenuated total reflectance Fourier transform spectroscopy (ATR-FTIR) for the binder recovered from glass fibre (S14) (black), and a reference library spectrum for PVA (grey). Panel E shows estimation of the molecular weight of the binder extracted from glass fibre (S14) by DLS. Points and solid line show a standard curve generated using PVA of known MW. Dashed lines indicate the hydrodynamic radius of the PVA extracted from S14 (11nm) and the inferred MW (36 kDa). Points and error bars indicate the mean and range respectively of duplicate DLS measurements. Panel F shows 1H NMR spectra used to estimate the percentage hydrolysis of the binder extracted from glass fibre (S14). The upper spectrum (labelled S14) is that of the extract, with unknown percentage hydrolysis and hence an unknown percentage of monomers bearing acetyl groups. The five spectra below were obtained using standards prepared by proportionately mixing 80% and 100% hydrolysed PVA to achieve a range of acetyl group content ranging from <0.1% up to 20% (as per labels to left of panel). The X-axis indicates chemical shift measured in parts per million (ppm) and the Y-axis shows relative intensity.

For further characterisation, the binder was extracted in liquid and freeze dried. Fourier transform infrared (FTIR) spectroscopy of the extract provided a fingerprint spectrum, which matched closely with the expected spectrum of polyvinyl alcohol (PVA) (Figure 4D) ^34, 35^.

The solubility and other properties of PVA vary widely according to molecular weight (MW) and degree of hydrolysis, and so we sought to further characterise the presumed PVA extracted from S14. We used dynamic light scattering (DLS) to compare the hydrodynamic radius of the PVA extract to those of PVA samples of known MW. The results were consistent with a MW in the range of 35 kDa (Figure 4E).

We then used 1H NMR to estimate the degree of hydrolysis of the polymer. The 1H NMR spectrum obtained (Figure 4F) was consistent with PVA, with a 2:1 ratio of the areas under the peaks at ∼1.6 ppm and 3.9 ppm (corresponding to hydrogens in the CH_2_ and CH environments respectively). Results obtained using a range of standards of varying percentage hydrolysis showed a clear relationship of the size of a peak at 2.05 ppm to the acetyl group content (the presence of which, in a sample of PVA, indicates incomplete hydrolysis). The spectra of the completely hydrolysed standard and the S14 extract were similar, with only a trace of a peak in this area, and so we concluded that the PVA extracted from S14 is likely to be completely hydrolysed.

### PVA enhances adenovirus stabilisation by SMT

Having identified PVA in the best-performing matrix, S14, we investigated whether the PVA may function not only as a binder but might actually contribute to vaccine thermostabilisation.

We reasoned that treatment with PVA might enhance the thermostabilisation performance of relatively poorly-thermostabilising matrices. We observed that application of PVA extracted from glass fibre (S14) significantly improved thermostabilisation performance of two of the three tested matrices, as assessed by infectious virus recovery after a four week 45 °C thermochallenge. Recovery from the polyester matrix (Leukosorb) was enhanced by 1.3 log_10_-fold, while enhancement from the cellulose matrix (ET CHR) was enhanced by 2 log_10_-fold (Figure 5A). Addition of PVA to a glass fibre matrix (Pall’s conjugate pad) was not beneficial. This matrix already contains a binder (Fig 2F), possibly PVA.

**Figure 5:**
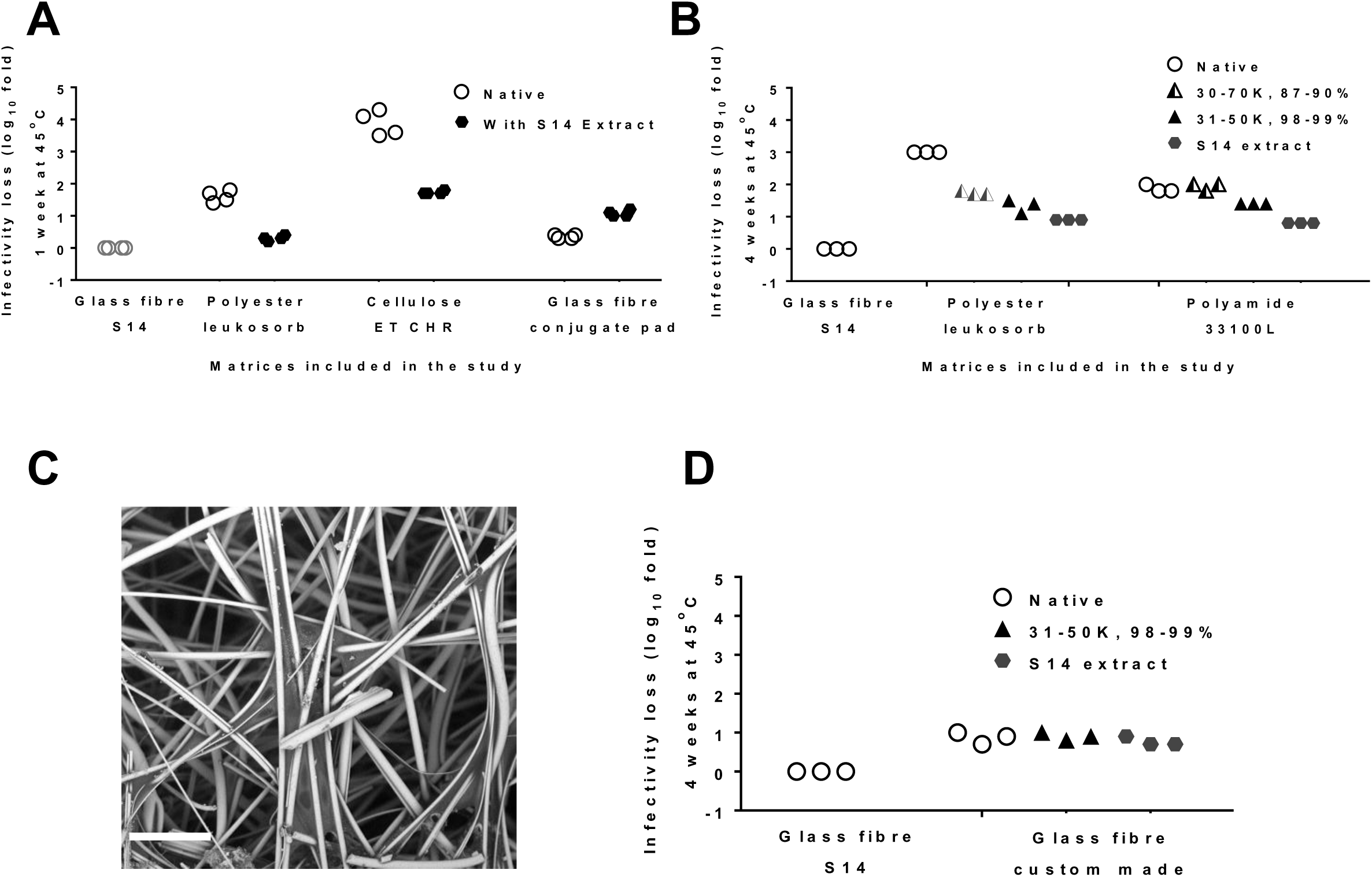
PVA improves vaccine thermostability on matrices. A PVA-containing extract prepared from S14 was dried onto various fresh matrices, followed by drying of vaccine in trehalose-sucrose buffer onto the treated matrices and untreated comparator matrices. Viable virus recovery was assessed after thermochallenge as indicated for each panel. S14 was included as a comparator in each experiment. Panel A shows loss in infectivity titre of vaccine in the absence (open circles) or presence (solid circles) of PVA extracted from glass fibre (S14), as compared to −80°C stored liquid control, after one week at 45 °C. Points indicate individual samples (four replicates per condition). Panel B shows difference in loss in infectivity titre of vaccine loaded in polyester (leukosorb) and polyamide (33100L) matrix modified by deposition of three different types of PVA prior to application and drying of vaccine. Dried samples were subjected to thermochallenge for four weeks at 45 °C. Points indicate individual samples (three replicates per condition). Panel C shows a scanning electron microscope image of the custom-made glass fibre matrix. Scale bar shows 50 µm. Panel D compares loss in infectivity titre of vaccine post-thermochallenge between standard 14 and custom-made glass fibre matrix modified by deposition of two different types of PVA prior to application and drying of vaccine. Dried samples were subjected to thermochallenge for four weeks at 45°C. Points indicate individual samples (three replicates per condition).

We proceeded to test whether the degree of hydrolysis of PVA had any impact on vaccine thermostability at 45°C for an extended period of 28 days. We tested polyester (Leukosorb), polyamide (33100L) and a custom-made glass fibre-based matrix after treatment with PVA extracted from S14 and two other commercially sourced polymers (Sigma) which were similar in size but differed in degree of hydrolysis (30-70 KDa / 87-90% hydrolysed, and 31-50 KDa/ >99% hydrolysed). We again observed substantial improvements in thermostabilisation performance, exceeding a 1 log_10_-fold increase in infectious virus recovery after thermochallenge, when PVA was applied to matrices which did not contain PVA at baseline (polyamide and polyester) (Figure 5B). The greatest enhancement was seen with the S14 extract, followed by the fully hydrolysed PVA, with the least enhancement seen with the incompletely hydrolysed PVA (Figure 5B).

### Towards development of custom nonwoven fabrics for vaccine thermostabilisation

Having characterised the geometry and composition of S14, and the contribution of PVA to its thermostabilisation performance, we proceeded to develop a bespoke matrix ‘in-house’. Although our ultimate aim is to produce biocompatible matrices, we sought as an initial step to test whether the understanding we had gained would enable us to produce an ‘in-house’ wet-laid glass fibre matrix which could replicate S14’s thermostabilisation performance. The new matrix had structural properties similar to those of S14 (area density 50.6±1.5 g.m^-2^, thickness 0.56±0.02 mm, mean flow pores 25±2.3 µm, absorption capacity 11.1±0.3 g/g, composed of borosilicate glass fibres with diameter 3.6±1.8 and length 1.3±0.6 mm; see table 2 for data relating to S14), and a similar appearance (Figure 5C). Although thermostabilisation performance of the custom-made matrix did not exactly match that of S14 (Figure 5D), it was closer than had previously been achieved with any of the other tested matrices (Fig 5B). As expected, further modification of the matrix with additional PVA had no effect.

## Discussion

The starting point for the present study was the observation, in our previous work, that the non-biocompatible glass-based S14 matrix out-performed a polypropylene matrix ^26^. This posed the question of which properties of the matrix could be relevant for SMT performance, and whether a more suitable matrix than S14 could be identified for clinical translation.

Our initial attempts to find a suitable commercially available matrix yielded disappointing results. Adenovirus stability on the tested matrices was poor (table 1). Multiple variables differed between each matrix, and we were therefore unable to perform experiments to clearly isolate the effect of a single variable. There was not a clear relationship between characteristics of the matrices and their stabilisation performance (Figures 2 and 3), with the possible exception of high enthalpic recovery of sugar glass (which is not a parameter which can readily be ‘designed in’ to a new matrix).

Our ability to use analysis of commercially available matrices to draw conclusions to guide design of new biocompatible matrices was thus limited. We therefore changed our strategy, seeking to characterise S14 in detail and produce similar matrices ‘in-house’, allowing us to identify features of S14 contributing to its stabilising performance.

Most significantly we found that PVA, present on the S14 matrix as a binder, appears to contribute to the stability of adenovirus (Figures 4 and 5). It was shown that fully hydrolysed PVA, similar to that we extracted from S14, was most beneficial in thermostabilisation (Figure 5). Polyvinyl alcohol is potentially suitable for use as an excipient in vaccine formulations: it is ‘generally regarded as safe’ (GRAS) and is also a FDA approved inactive ingredient for parenteral use ^36^. PVA has previously been explored as an excipient in a number of studies of bio-macromolecular stability. It has been found to be beneficial in some formulations of proteins, including insulin ^37^, but benefit has not been seen consistently in other studies ^38 39 40^. PVA may contribute to protein stability by hydrogen bonding of hydroxyl groups in PVA to the proteins. In dried protein formulations, PVA has also been shown to prevent deamidation more potently than another widely used polymeric excipient, polyvinyl pyrrolidone ^41^.

We now intend to develop the SMT method towards clinical application. There are two principal obstacles to this goal: robust and GMP compliant execution of the process, and the availability of a suitable matrix for GMP production. Robustness and GMP compliance of the process are clearly interlinked. We believe several of the challenges of GMP execution of the SMT process can be addressed by execution of drying within a lyophilizer without freezing or the application of vacuum: existing large-scale GMP lyophilisation facilities could provide the necessary controlled temperature, low humidity, aseptic environment. With respect to robustness, we have found in recent work that the stability achieved by the process can be highly sensitive to deviations from the intended conditions and, more troublingly, some unexplained inconsistency in performance can occur: work is ongoing to address these issues. With respect to the matrix, GMP execution is likely to require development of a new non-woven. In addition to replicating the stabilising performance of S14, such a matrix needs to be biocompatible, to have good mechanical integrity (in particular, without shedding fibres into the reconstituted product), and to be produced in line with the quality requirements for a GMP raw material. We are now using the data provided by the present study in order to develop such a matrix, mimicking the physical properties of S14 and making using of the beneficial effect of PVA, but without the problematic use of glass.

## Acknowledgments

We are grateful for the assistance of the Jenner Institute Viral Vector Core Facility in the production and immunostaining of adenoviruses, to Pall Corporation, GE Healthcare and Hollingsworth-Vose for providing filter media and technical discussion and to Dr. Isaac Rubens Martínez Pardo from RMIT University, Australia for his assistance in FTIR data analysis

